# Developing Human Pluripotent Stem Cell-Based Cerebral Organoids with a Controllable Microglia Ratio for Modeling Brain Development and Pathology

**DOI:** 10.1101/2020.10.09.331710

**Authors:** Ranjie Xu, Andrew J. Boreland, Xiaoxi Li, Caroline Erickson, Mengmeng Jin, Colm Atkins, Zhiping Pang, Brian P. Daniels, Peng Jiang

## Abstract

Microglia, as brain-resident macrophages, play critical roles in brain development, homeostasis, and disease. Microglia in animal models cannot accurately model the properties of human microglia due to notable transcriptomic and functional differences between human and other animal microglia. Efficient generation of microglia from human pluripotent stem cells (hPSCs) provides unprecedented opportunities to study the function and behavior of human microglia. Particularly, incorporating hPSCs-derived microglia into brain organoids facilitates their development in a 3-dimensional context, mimicking the brain environment. However, an optimized method that integrates an appropriate amount of microglia into brain organoids at a proper time point, resembling in vivo brain development, is still lacking. Here, we report the development of a new brain region-specific, microglia-containing organoid model by co-culturing hPSCs-derived primitive neural progenitor cells (pNPCs) and primitive macrophage progenitors (PMPs). In these organoids, hPSCs-derived pNPCs and PMPs interact with each other and develop into functional neurons, astroglia, and microglia, respectively. Importantly, the numbers of human microglia in the organoids can be controlled, resulting in a cell type ratio similar to that seen in the human brain. Using super-resolution microscopy, we demonstrate that these human microglia are able to phagocytize neural progenitor cells (NPCs) and apoptotic cells, as well as to prune synapses at different developmental stages of the organoids. Furthermore, these human microglia respond to Zika virus infection of the organoids, as indicated by amoeboid-like morphology, increased expression of gene transcripts encoding inflammatory cytokines, and excessive pruning of synaptic materials. Together, our findings establish a new microglia-containing brain organoid model that will serve to study human microglial function in a variety of neurological disorders.

## Introduction

Microglia, as the innate immune cells of the central nervous system (CNS), play critical roles in neurodevelopment and tissue homeostasis. During brain development, microglia phagocytize apoptotic cells, prune redundant synapses, and regulate neurogenesis and axonal growth (Casano and Peri, 2015; Wolf et al., 2017). Under homeostatic conditions, microglia survey the CNS environment, detect and respond to protein aggregates, pathogens, and other damage that may endanger the CNS and compromise tissue homeostasis (Bahmad et al., 2018; Casano and Peri, 2015). Increasing evidence has also demonstrated that microglial dysfunction is implicated in a broad range of neurological disorders, including schizophrenia, Alzheimer’s disease, and traumatic brain injury (Wolf et al., 2017). Moreover, genome-wide association studies show that many neurological disease risk genes are highly, and sometimes exclusively, expressed in microglia (Gosselin et al., 2017). Thus, there is a compelling incentive to understand the role of microglia in brain development and disease.

Most of our knowledge of microglia has relied on animal microglia, such as those derived from rodents or zebrafish (Pocock and Piers, 2018). However, human microglia are distinct from microglia in other animals, including differential expression of the complement cascade, phagocytic functions, and expression of susceptibility genes to neurodegeneration (Galatro et al., 2017; Geirsdottir et al., 2020; Gosselin et al., 2017). Because of these innate species differences, there is an urgent need to develop more representative strategies for investigating the unique functions of human microglia in health and disease. Recent advances in stem cell technologies have enabled the generation of human microglia from human pluripotent stem cells (hPSCs), greatly facilitating the study of the pathophysiology of human microglia (Douvaras et al., 2017; Guttikonda et al., 2021; Haenseler et al., 2017; McQuade et al., 2018; Muffat et al., 2016).

Notably, previous studies demonstrated that the transcriptomic profile and morphology of hPSC-derived microglia cultured under 2-dimensinoal (2D) conditions are different than microglia in the *in vivo* brain. This is in part due to limited cell-cell and cell-matrix interactions in the 2D cultures(Gosselin et al., 2017; Jiang et al., 2020). Brain organoids derived from hPSCs in a 3D context contain diverse neural cell types, recapitulate embryonic brain architecture, and provide unprecedented opportunities to study human brain development and the pathogenesis of neurodevelopmental disorders (Di Lullo and Kriegstein, 2017; Qian et al., 2020). Thus, organoids may enhance human microglial survival and functional development, since human microglia in organoids likely are exposed to brain region-specific signaling cues and interact with other cell types, including neurons and astrocytes. Previous studies have demonstrated the feasibility of generating microglia-containing organoids (Abud et al., 2017; Ormel et al., 2018). However, these models have significant limitations. Abud et al. reported integration of mature microglia into cerebral organoids that had been cultured for a long time period (Abud et al., 2017). During early embryonic development, primitive macrophage progenitors (PMPs) originating from the yolk sac migrate to the CNS through the circulation (Ginhoux et al., 2010; Ginhoux and Guilliams, 2016). Those PMPs and neural progenitor cells (NPCs) interact with each other and undergo differentiation and maturation together *in vivo* (Li and Barres, 2018). As such, the model reported by Abud et al. is limited in examining the interactions between microglia and NPCs or developing neurons because microglia are introduced into the organoids at a stage when neurons have already developed. The timing of integrating microglia into organoids may impact neuronal population and maturation (Del Dosso et al., 2020). Recently, in order to promote mesoderm differentiation in brain organoids, Ormel et al. developed methods to generate organoids without using dual SMAD inhibition to induce neural differentiation (Ormel et al., 2018). Microglia innately develop in these cerebral organoids, simultaneously with neural differentiation. However, due to the absence of SMAD inhibition of direct cell differentiation, the heterogeneity of cell types in those organoids is high and uncontrollable.

In this study, we hypothesize that by co-culturing hPSCs-derived primitive neural progenitor cells (pNPCs) and PMPs, we can generate developmentally appropriate and brain region-specific microglia-containing brain organoids. The fact that the starting numbers of PMPs and pNPCs can be readily and precisely controlled further reduces the variability between individual organoids. Importantly, the numbers of human microglia in the organoids can be controlled, resulting in a cell type ratio similar to that seen in the brain. Moreover, we demonstrate that in our microglia-containing organoids, human microglia exhibit phagocytic activity and synaptic pruning function. We also show that human microglia are dynamic in response to Zika virus (ZIKV) infection and find that ZIKV causes inflammation and excessive elimination of synaptic components by microglia.

## Results

### Generation of brain organoid with a controllable microglial ratio

During embryonic CNS development, yolk sac-derived myeloid progenitors migrate into the developing neural tube, where both NPCs and myeloid progenitors interact, proliferate, and mature into neurons, astrocytes, oligodendrocytes, and microglia (Colonna and Butovsky, 2017). To mimic this early neurodevelopmental process, we designed a strategy to induce NPCs and PMPs from hPSCs first, and then co-culture NPCs and PMPs in 3D to generate microglia-containing brain organoids (Figure 1A). To induce neural differentiation, we used three hPSC lines, including two healthy individual-derived human induced pluripotent stem cells (hiPSCs) and one human embryonic stem cell (hESC), with dual inhibition of SMAD signaling(Chambers et al., 2009). At day 7, we plated the embryoid bodies (EB) on dishes, and on day 14, we manually separated neural rosettes from surrounding cells and further expanded as pNPCs, as described in our previous studies (Chen et al., 2016; Xu et al., 2019). The identity of hPSCs-derived pNPCs was confirmed by staining with Nestin, a marker for NPCs (Wiese et al., 2004) (Figure 1B). To induce microglia differentiation and obtain PMPs, we followed a published protocol (Haenseler et al., 2017; Xu et al., 2020). At day 5, yolk sac embryoid bodies (YS-EB) were plated on dishes, and PMPs were released from attached YS-EB into the supernatant starting around day 20; PMPs were continuously produced for more than three months, with a cumulative yield about 40-fold higher than the number of the input hPSCs. The identity of hPSCs-derived PMPs was confirmed by staining with CD235, a marker for YS primitive hematopoietic progenitors (Claes et al., 2018), and CD43, a marker for hematopoietic progenitor-like cells (Claes et al., 2018). Moreover, the PMPs were highly proliferative, as indicated by Ki67 staining (Figure 1C). Next, we cultured pNPCs and PMPs together and supplemented with mitogen FGF-2 for another two weeks (up to day 35), which is termed as the proliferation stage. At day 35, we transitioned organoids to neural differentiation medium, which contained growth factors, such as BDNF, GDNF to promote neural maturation, and IL-34, GM-CSF, which are important for microglia survival and maturation (Haenseler et al., 2017). We termed this stage as the neural differentiation stage. Organoids can be maintained in this stage for a long period for further characterization.

**Figure 1.**
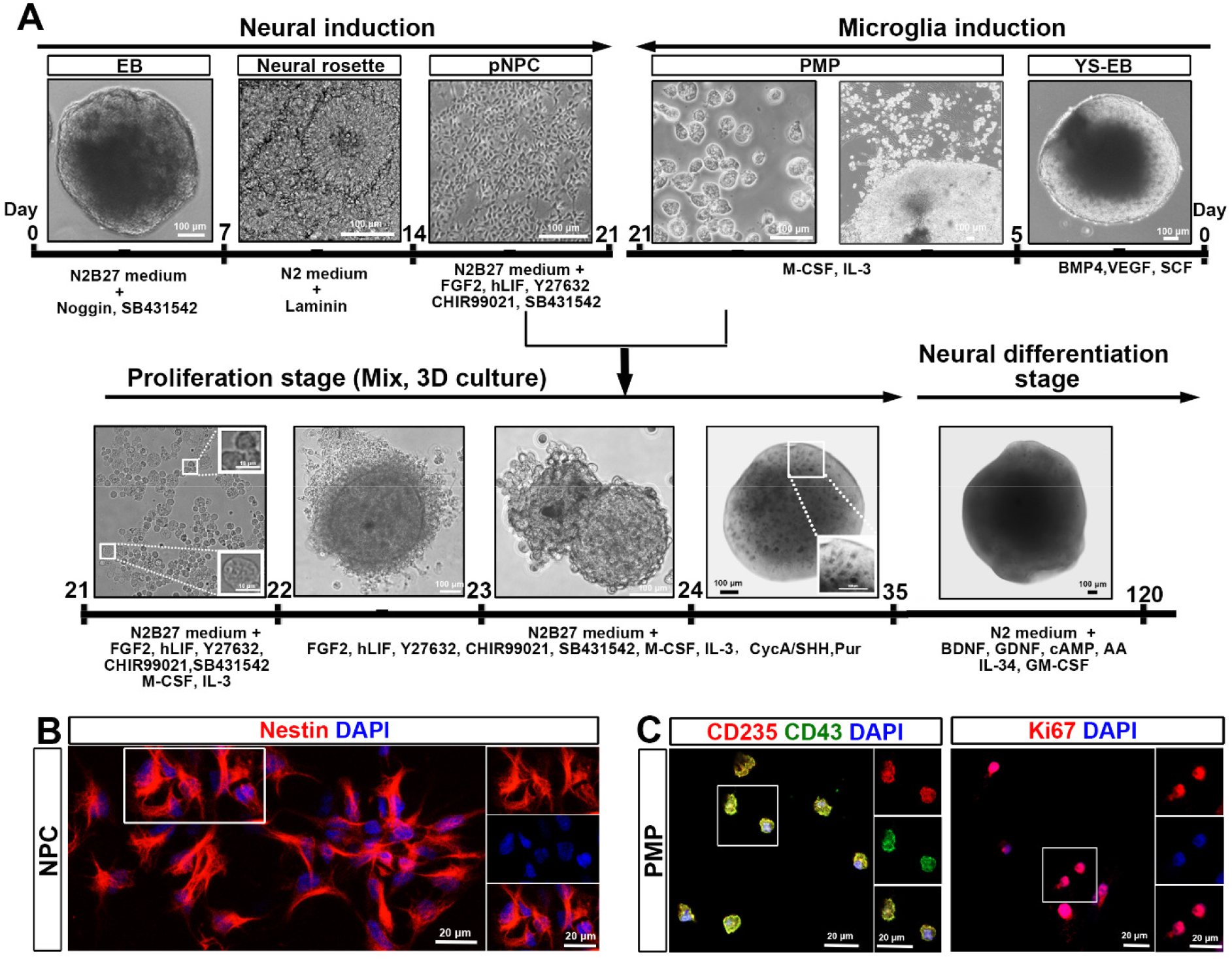
Generation of microglia-containing brain organoids. (A) schematic procedure for deriving microglia-containing brain organoids by co-culture of primitive neural progenitor cells (pNPCs) and primitive macrophage progenitors (PMPs) under 3-dimensional (3D) conditions. Scale bars: 100 μm or 10 μm in the original or enlarged images, respectively. (B) Representatives of Nestin^+^ pNPC. Scale bars: 20 μm in the original or enlarged images. (C) Representatives of CD235^+^, CD43^+^, and Ki67^+^ PMPs. Scale bars: 20 μm in the original or enlarged images.

### PMPs develop into microglia in brain organoids

Microglia account for around 5-10% of total CNS cells (Frost and Schafer, 2016). To mimic the percentage of microglia in the CNS, we tested two different starting ratios of pNPCs and PMPs: 9:1 or 7:3. We found that starting with 7000 pNPCs and 3000 PMPs generated organoids with around 8% microglia in the total cell population (Figure S1A and S1B). Also, organoids gradually increased in size during culture (Figure S1C). To confirm whether PMPs mature into microglia in the brain organoid, we examined the identity of microglia at different stages. At the proliferation stage (day 35), around 13% (13.7% ± 4.6) of cells in the organoid expressed PU.1, a transcription factor that is necessary for microglial differentiation (McKercher et al., 1996), and the majority of cells in the organoid were proliferative, as indicated by Ki67 staining (Figures 2A and 2G). Among all the cells, about 5% (4.6% ± 1.5) of cells expressed both PU.1 and Ki67, suggesting PMPs started to differentiate into microglia and were proliferative at this stage (Figure 2A). Next, we examined the identity of microglia at neural differentiation stage by using a panel of microglia markers. At day 45, organoids co-expressed Iba1, a canonical macrophage/microglia marker (Butovsky and Weiner, 2018), and CD45, which is expressed by all nucleated hematopoietic cells (Ford et al., 1995) (Figure 2B). At day 50, appropriate microglia morphology became increasingly evident. About 8% (8.0 % ± 1.6) of cells expressed Iba1, and most of these Iba1-expressing microglia showed ramified morphology (Figures 2C and 2H). Besides, some of the cells in day 55 organoids also co-expressed CD11b and CD45, as well as Iba1 and PU.1 (Figures 2D and 2E). We also found cells that expressed TMEM119 (Figure 2F), a marker that distinguishes microglia from other macrophages (Satoh et al., 2016). Taken together, these results demonstrate that PMPs differentiate into microglia in our brain organoids.

**Figure 2.**
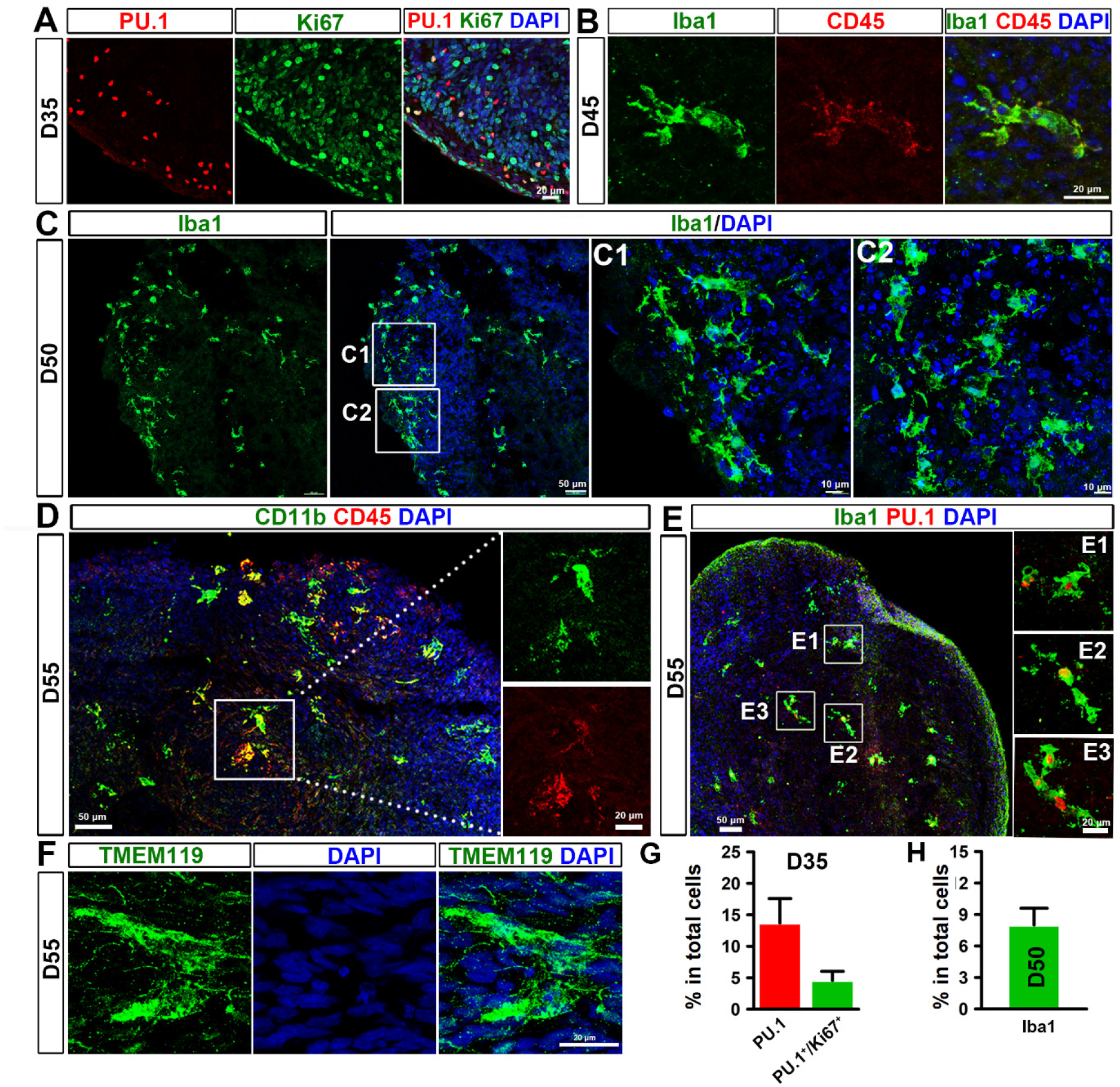
Differentiation of PMPs into microglia in brain organoids. (A) Representatives of PU.1^+^/Ki67^+^ cells in day 35 brain organoids. Scale bars: 20 μm. (B) Representatives of Iba1^+^/CD45^+^ cells in day 45 brain organoids. Scale bars: 20 μm. (C) Representatives of Iba1^+^ cells in day 50 brain organoids. Scale bars: 50 μm or 10 μm in the original or enlarged images, respectively. (D) Representatives of CD11b^+^/CD45^+^ cells in day 55 brain organoids. Scale bars: 50 μm or 20 μm in the original or enlarged images, respectively. (E) Representatives of Iba1^+^/PU.1^+^ cells in day 55 brain organoids. Scale bars: 50 μm or 20 μm in the original or enlarged images, respectively. (F) Representatives of TMEM119^+^ cells in day 55 brain organoids. Scale bars: 20 μm. (G) Quantification of pooled data from two hiPSC lines and one hESC line showing the percentage of PU.1^+^, and PU.1^+^/Ki67^+^ cells in day 35 organoids (n = 3). Three independent experiments were performed and for each experiment, 4-6 organoids from each hPSC line were used. Data are presented as mean ± s.e.m. (H) Quantification of pooled data from hiPSC and hESC lines showing the percentage of Iba1^+^cells in day 50 organoids (n = 3). Three independent experiments were performed, and for each experiment, 4-6 organoids from each hPSC line were used. Data are presented as mean ± s.e.m.

### Neural maturation in microglia-containing brain organoid

We then examined neural maturation in microglia-containing brain organoids. Previous studies demonstrate microglia can modulate neurogenesis by providing neurotrophic factors and cytokines, such as IGF1 and IL-1β (Casano and Peri, 2015; Shigemoto-Mogami et al., 2014; Ueno et al., 2013). Thus, we examined the mRNA expression of *IGF1* and *IL-1β* in organoids with or without microglia at the early proliferation stage. At day 27 when pNPCs were cultured alone or co-cultured with PMPs for 7 days, we found that the mRNA expression of *IGF1* and *IL-1β* dramatically increased in the organoids containing microglia, as compared to the organoids without microglia (around 1757-fold for *IGF1* and 242-fold for *IL-1β*; Figure S1D). These results suggest that microglial cells in the organoids might promote neurogenesis from NPCs by providing these factors. At day 40, ventricular zone-like regions were seen in organoids, which contained Ki67^+^ and SOX2^+^ progenitors, as well as TUJ-1^+^ immature neurons (Figure 3A). At day 55, organoids had a large number of immature neurons, as indicated by the expression of Doublecortin (Dcx), a marker of immature neurons (Gleeson et al., 1999) (Figure 3B). As shown in Figure 3C and 3D, NeuN^+^ neurons and S100β^+^ astrocytes were detected in day 55 and 75 organoids. To further confirm whether neurons were functionally mature in the microglia-containing organoids, we performed whole-cell patch-clamp recordings of neurons from acutely sectioned organoid slices (Figure 3E). Neurons recorded were simultaneously loaded with Biotin Ethylenediamine HBR for morphological representation and displayed complex arborization, long-distance projections, and spine-like structures (Figure 3F). We found that human neurons cultured in the 3D microglia-containing organoids were functionally mature, as indicated by these neurons robustly expressing voltage-gated K^+^ and Na^+^ currents (Figure 3G), firing spontaneous action potentials (Figure 3H) and repetitive evoked action potentials upon step-wise current injections (Figure 3I). More importantly, neurons in the organoids (90 days in culture) exhibit spontaneous post-synaptic currents (PSCs) (Figure 3J). Taken together, these results show that pNPCs differentiate into functional neurons and astrocytes in the presence of microglia in our organoids.

**Figure 3.**
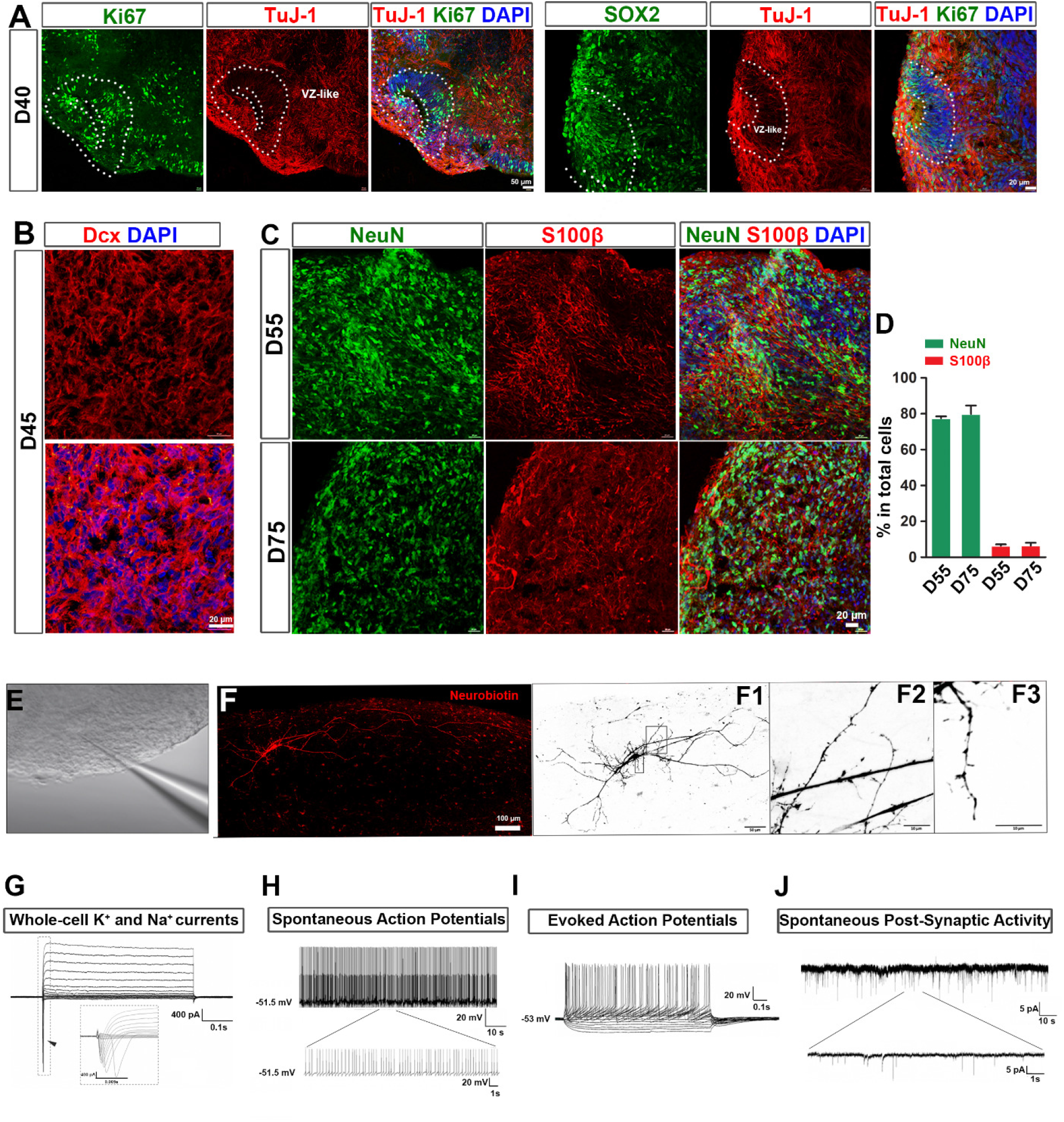
Neural maturation in microglia-containing brain organoid. (A) Representatives of Ki67^+^, SOX2^+^, and Tuj1^+^ cells in day 40 microglia-containing brain organoids. Scale bars: 50 μm or 20 μm as indicated. (B) Representatives of DCX^+^ cells in day 45 microglia-containing brain organoid. Scale bars: 20 μm. (C) Representatives of NeuN^+^ and S100β^+^ cells in day 55 or 75 brain organoids with or without microglia. Scale bars: 20 μm. (D) Quantification of pooled data from two hiPSC lines and one hESC line showing the diameter of day 35, 55 or 75 brain organoids with or without microglia. (n = 3), Three independent experiments were performed and for each experiment, 4-6 organoids from each hPSC line were used. Student’s *t* test. *NS* represents no significance. Data are presented as mean ± s.e.m. (E) A representative image showing a patched neuron in acute organoid slice. (F) Sample images of a neuron filled with Biotin Ethylenediamine HBR from a day 92 microglia-containing organoid after whole-cell recording. Complex arborization (F1) and spine-like structures (F2, F3) were identified. The images are inverted to a white background to help visualize neuron morphology. (G) Representative traces of whole-cell currents recorded from neurons in acutely sectioned organoid (90 day old in culture) slices. Insert: fast activation/inactivation voltage dependent sodium currents. (H) Representative traces of spontaneous action potentials recorded from neurons in organoid (90 day old in culture) slices. (I) Example of repetitive action potentials evoked by stepwise currents injections. (J) Representative traces of spontaneous postsynaptic currents (PSCs) recorded from neurons in sliced collected from a day 92 organoid.

### Microglia phagocytize NPCs and remove apoptotic cells in brain-region specific organoids

As the resident macrophages of the CNS, microglia play vital roles in neurodevelopment and tissue homeostasis mediated in part by their phagocytic functions. Microglia phagocytize NPCs to control their relative abundance during development, clear extracellular debris and remove dead cells to maintain CNS homeostasis (Wolf et al., 2017). To investigate whether microglia have phagocytic functions in organoids, we examined the expression of CD68, a marker for the phagolysosome (Weinhard et al., 2018). We found the vast majority of microglia expressed CD68 (Figures 4A and 4B, and S2A), suggesting their active phagocytic functions. Next, we patterned the organoids into brain region-specific organoids, including the dorsal forebrain, as indicated by dorsal forebrain marker PAX6 (Sussel et al., 1999), or ventral forebrain, as indicated by ventral forebrain marker NKX2.1 (Sussel et al., 1999). As shown in Figures 4C and 4D, we observed that around 13% (12.9 ± 4.0%) or 15% (15.4 ± 2.5%) of CD68^+^ microglia phagocytized PAX6^+^ or NKX2.1^+^ NPCs in day 45 dorsal or ventral forebrain organoids, respectively (Figures 4C,4D,4G, and 4H). Additionally, CD68^+^ microglia phagocytized developing neurons, as indicated by the intermediate progenitor stage marker Tbr2 (Sessa et al., 2010) (Figures 4E, S2B). We found around 18% (18.4 ± 5.6%) of CD68^+^ microglia phagocytized Tbr2^+^ developing neurons in day 55 organoids (Figure 4I). To confirm whether microglia can remove apoptotic cells, we co-stained CD68 and activated Caspase-3, an apoptotic cell marker, and found that around 15% (15.0 ± 0.8%) of CD68^+^ microglia phagocytized the apoptotic cells (Figures 4F, 4J, S2C, S2D). Occasionally, we also found that microglia responded and aggregated around foreign bodies in the organoid caused by embedded fragments from a glass pipette tip (Figure S2E).

**Figure 4.**
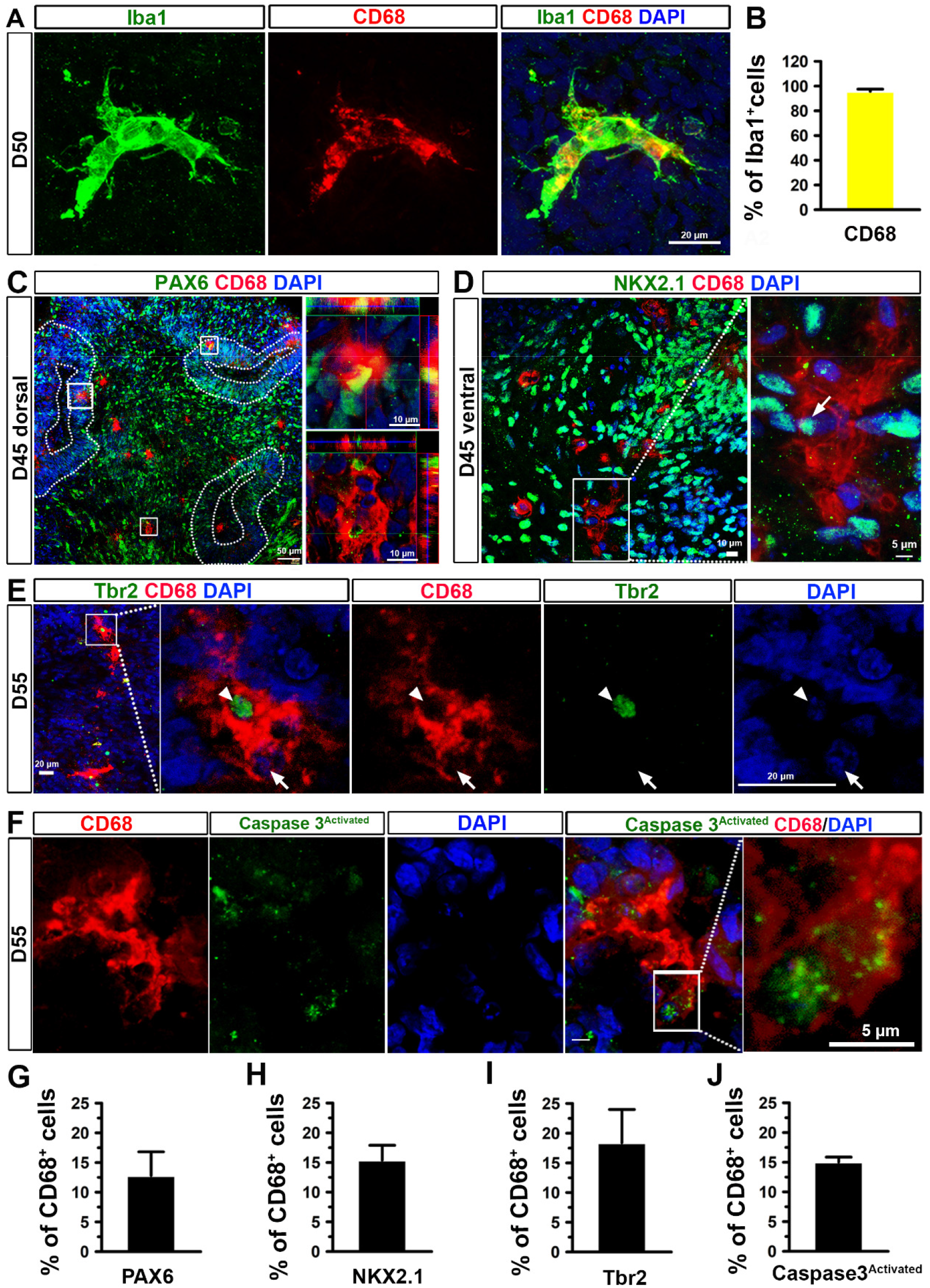
Phagocytic function of microglia in brain-region specific organoids. (A) Representatives of Iba1^+^/CD68^+^ cells in day 50 brain organoids. Scale bars: 20 μm. (B) Quantification of pooled data from two hiPSC lines and one hESC line showing the percentage of CD68^+^ cells in total Iba1^+^ cells in day 50 organoids (n = 3, three lines each time, 4-6 organoids each line each time). Data are presented as mean ± s.e.m. (C) Representative images showing CD68^+^ microglia phagocytosing PAX6^+^ dorsal forebrain NPCs in day 46 dorsal forebrain organoids (enlarged images). Scale bars: 50 μm or 10 μm in the original or enlarged images, respectively. (D) Representative images showing CD68^+^ microglia phagocytosing NKX2.1^+^ ventral forebrain NPCs in day 46 ventral forebrain organoids (enlarged images). Scale bars: 10 μm or 5 μm in the original or enlarged images, respectively. (E) Representative images showing CD68^+^ microglia phagocytosing Tbr2^+^ intermediate progenitor cells in day 56 organoids (enlarged images). Scale bars: 10 μm or 5 μm in the original or enlarged images, respectively. (F) Representative images showing CD68^+^ microglia phagocytosing activated Caspase3^+^ apoptotic cells in day 56 organoids (enlarged images). Scale bars: 5 μm. (G and H) Quantification of pooled data from two hiPSC lines and one hESC line showing the percentage of PAX6^+^ (G) or NKX2.1^+^ cells (H) in total CD68^+^ cells in day 45 organoids (n = 3, three lines each time, 4-6 organoids each line each time). Data are presented as mean ± s.e.m. (I and J) Quantification of pooled data from two hiPSC lines and one hESC line showing the percentage of Tbr2^+^ (I) or activated Caspase 3^+^ cells (J) in total CD68^+^ cells in day 55 organoids (n = 3, three lines each time, 4-6 organoids each line each time). Data are presented as mean ± s.e.m.

### Microglia prune synapses in brain organoids

As shown in Figure 5A, Iba1^+^ microglia resided closely to MAP2^+^ neuronal dendrites. We also observed that synaptic puncta, labeled by PSD95, a post-synaptic marker, and synapsin 1, a pre-synaptic marker, as well as the juxtaposition of pre- and post-synaptic puncta complexes distributed along the MAP2^+^ dendrites (Figure 5B and S3A). We further quantified the density of synapsin 1^+^, PSD95^+^, and colocalized PSD95^+^ puncta (PSD95^+^ puncta in the juxtaposition of puncta complexes), indicated by the number of puncta per 10 μm^2^ MAP2^+^ dendrites, following the methods reported in previous studies (Catarino et al., 2013; Patzke et al., 2019) (Ciani et al., 2011). As shown in Figure 5C, in day 55 organoids, the density of synapsin 1^+^, PSD95^+^, and colocalized PSD95^+^ puncta is 5.4 ± 1.2, 3.7 ± 1.0, and 1.4 ± 0.5, respectively. However, the density of the juxtaposition of puncta was lower than that of single synapsin 1^+^ or PSD95^+^ puncta along with dendrites areas in day 55 organoids (Figure S3D). To examine whether these human microglia are able to prune synapses in our organoids. We triple-stained Iba1, CD68, and PSD95 and employed a super-resolution imaging technique to visualize synapse engulfment by microglia. As shown in Figure 5D, PSD95^+^ puncta were localized within CD68^+^ phagolysosomes in Iba1^+^ microglia, indicating that microglia phagocytized synaptic proteins in day 55 organoids. Similarly, synapsin I^+^ puncta were found within CD68^+^ phagolysosomes (Figure S3B). Next, we triple-stained CD68 with PSD95 and synapsin 1. The 3D reconstruction images clearly showed that PSD95^+^ and synapsin 1^+^ puncta were colocalized within CD68^+^ microglia, further indicating synaptic pruning function of the microglia in organoids (Figure 5E). In addition, 3D-surface rendered images show that single pre-, post-synaptic puncta, as well as the juxtaposition of puncta complexes, were inside of CD68^+^ phagolysosomes (Figure S3C). We quantified the number of synapsin 1^+^, PSD95^+^, and colocalized PSD95^+^ puncta per 100 μm^3^ CD68^+^ phagolysosomes (Figure 5F). The quantification result shows fewer colocalized PSD95^+^ puncta (0.5± 0.2) than single synapsin 1^+^ puncta (5.6 ± 1.3) or PSD95^+^ puncta (6.7 ± 0.8) inside CD68^+^ phagolysosomes.

**Figure 5.**
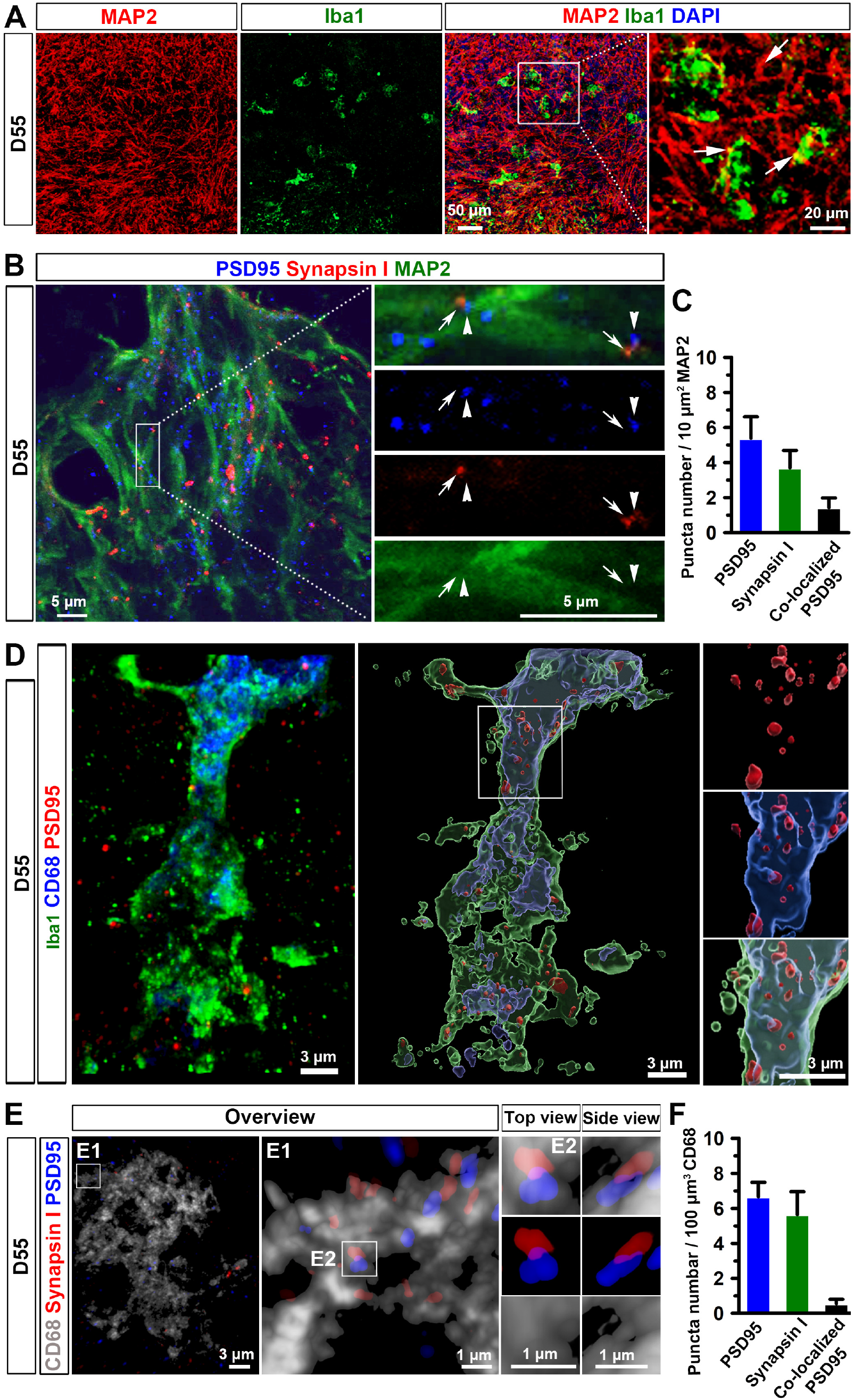
Microglia prune synapses in microglia-containing brain organoids. (A) Representative images showing Iba1^+^ microglia interact with MAP2^+^ neuronal processes in day 55 brain organoids. Scale bars: 50 μm or 20 μm in the original or enlarged images, respectively. (B) Representative images showing PSD95^+^ and synapsin 1^+^ synaptic puncta, and the juxtaposition of pre- and postsynaptic puncta complexes along MAP2^+^ processes in day 55 brain organoids. Arrows and arrowheads indicate the juxtaposition of pre- and postsynaptic puncta, respectively. Scale bars: 5 μm in the original or enlarged images. (C) Quantification of pooled data from two hiPSC lines and one hESC line showing the density of synapsin 1^+^, PSD95^+^, and colocalized PSD95^+^ per 10 μm^2^ MAP2 in day 55 organoids (n = 3, three lines each time, 4-6 organoids each line each time). Data are presented as mean ± s.e.m. (D) Representative images showing co-localization of Iba1^+^, CD68^+^, and PSD95^+^ staining in day 55 organoids. Left, a raw fluorescent super-resolution image. Right, a 3D-surface rendered image from the same raw image. Scale bars, 3 μm. (E) Representative 3D reconstruction of super-resolution images showing CD68^+^ microglial engulfment of synapsin 1 and PSD95 in day 55 organoids. Scale bars, 3 and 1 μm in the original or enlarged images, respectively. (F) Quantification of pooled data from two hiPSC lines and one hESC line showing the number of synapsin 1^+^, PSD95^+^, and colocalized PSD95^+^ per 100 μm^3^ CD68^+^ microglial phagolysosomes in day 55 organoids (n = 3, three lines each time, 4-6 organoids each line each time). Data are presented as mean ± s.e.m.

### Microglia respond to Zika virus (ZIKV) infection in brain organoid

We further explored whether our organoid model can be employed to study microglial functions under diseased or pathological conditions. Brain organoids have been used to model ZIKV infection before (Abreu et al., 2018; Dang et al., 2016; Garcez et al., 2016; Qian et al., 2016). However, microglia were missing in most of the previous organoid models and how human microglia are involved in ZIKV-related neuropathology is largely unknown. To address this question, we infected our microglia-containing organoids with two different stains of ZIKV, MR766 and Fortaleza. Consistent with previous studies, ZIKV-MR776 exhibited greater virulence than Fortaleza (Esser-Nobis et al., 2019), as indicated by the significant morphological abnormalities and cell detachment in the organoid (Figure 6A, S4A). Thus, we chose to use the MR766 strain for the following experiments. We measured viral replication using a standard plaque assay, which confirmed productive infection of the organoid (Figure 6B). Furthermore, flavivirus specific envelope (E)-protein, was found to be highly expressed in ZIKV-exposed organoids but not in mock-infected organoids, further indicating the success of ZIKV infection (Figure 6C). Interestingly, microglial morphology dramatically changed after virus exposure and became more amoeboid-like, indicated by the significantly decreased number of endpoints and process length, suggesting microglial activation (Figures 6D, 6E). In addition, we examined GFAP expression in organoids after ZIKA infection. As shown in Figures S4B and S4C, the percentage of GFAP^+^ cells in organoids significantly increased after ZKAV exposure compared with mock-infected control, suggesting astrocytes respond to ZIKV challenge, consistent with previous studies (Buttner et al., 2019; Huang et al., 2018). Next, we examined the mRNA expression of *AXL*, a potential entry receptor for ZIKV, which is highly expressed in human astrocyte and microglia (Di Lullo and Kriegstein, 2017; Meertens et al., 2017; Qian et al., 2016) and critical for infection of these human glial cells, but not human NPCs (Di Lullo and Kriegstein, 2017; Meertens et al., 2017). Interestingly, we found that mRNA expression of *AXL* significantly upregulated after ZIKA infection (Figure S4D). We further examined AXL protein levels in organoids by immunostaining and found increased AXL expression after ZIKA infection (Figure S4E, S4F). To confirm whether microglia and/or astrocytes contribute to the increased level of AXL, we stained AXL with CD68 and GFAP. We observed significantly increased expression of AXL in CD68^+^ microglia after ZIKA infection. Few GFAP^+^ astrocytes expressed AXL and there was no significant differences in the expression of AXL in GFAP^+^ astrocytes between mock and ZIKV infection groups (Figure S4E, S4F). These results suggest that microglial, rather than astroglial, increase of AXL expression contributes to the AXL upregulation after ZIKA infection in organoids. We then examined the expression of inflammatory cytokines *IL-6*, *IL-1β*, and *TNF-α* by qRT-PCR. Expression of these inflammatory cytokines was significantly increased in the ZIKV-exposed organoids, consistent with previous studies (Abreu et al., 2018; Figueiredo et al., 2019)(Figure 6F). Type I interferon signaling is involved in response to viral and bacterial infections (Ivashkiv and Donlin, 2014). We examined the mRNA expression of *INFAR1* and *INFAR2* and found that both *INFAR1* and *INFAR2* mRNA expression increased after the ZIKV exposure (Figure 6G). These data confirm successful infection of the organoid, as well as the establishment of a robust innate immune response.

**Figure 6.**
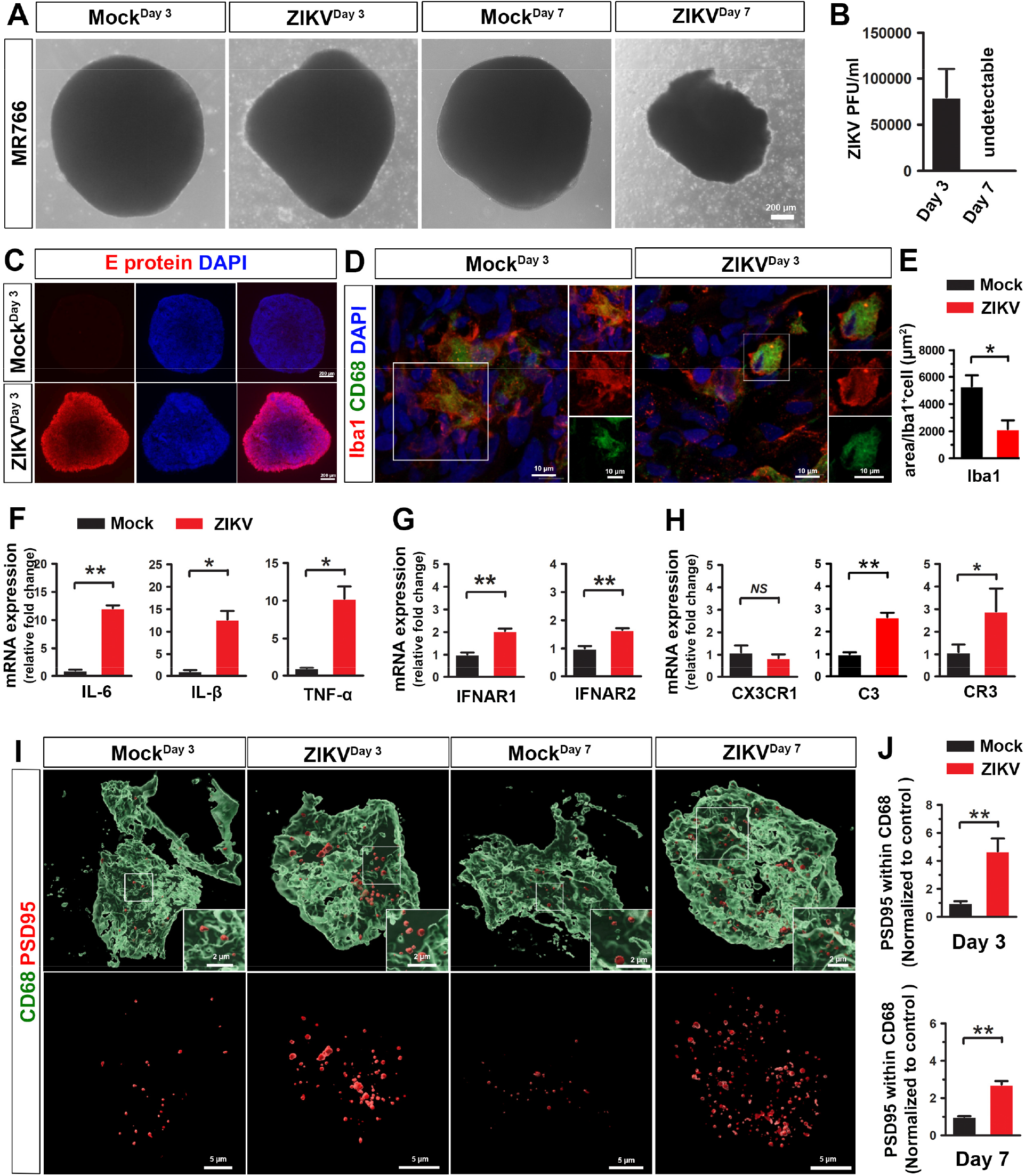
Responses of microglia to ZIKV infection in brain organoids. (A) Representative phase-contrast images showing the morphology of day 75 microglia-containing organoid after three and seven days of ZIKV or mock infection. Scale bars: 200 μm. (B) Quantitation of infectious ZIKV particles in the culture medium by plaque assay. (C) Representative images showing E protein expression in day 75 organoids after three days of ZIKV or mock infection. Scale bars: 200 μm. (D) Representatives of Iba1^+^/CD68^+^ microglia in day 75 organoids after three days of ZIKV or mock infection. Scale bars: 50 μm or 20 μm in the original or enlarged images, respectively. (E) Quantification of the number of endpoints and process length of Iba1^+^ microglia in day 75 organoids after three days of ZIKV or mock infection (n = 3, three independent times infections, each time, 8-12 organoids from two hiPSC lines). Student’s *t* test. * represents *P < 0.01. Data are presented as mean ± s.e.m. (F) qPCR analysis of *IL-6*, *IL-β*, and TNF-α mRNA expression in day 75 organoid after three days of ZIKV or mock infection. (G) qPCR analysis of *IFNAR1* and *IFNAR2* mRNA expression in day 75 organoid after three days of ZIKV or mock infection. (H) qPCR analysis of *CX3CR1*, *C3*, and *CR3* mRNA expression in day 75 organoids after three days of ZIKV or mock infection. (I) Representative 3D reconstruction of super-resolution images showing PSD95^+^ puncta within CD68^+^ microglial lysosomes in day 75 organoids after three days of ZIKV or mock infection. Scale bars, 5 and 2 μm in the original or enlarged images, respectively. (J) Quantification of PSD95^+^ puncta engulfment in CD68^+^ microglial phagolysosomes in day 75 organoids after three days of ZIKV or mock infection. Notably, larger volumes of PSD95^+^ puncta were seen inside microglia in response to ZIKV, as compared to mock infection group. (n = 3, three independent times infections, each time, 9-12 organoids from two hiPSC lines and one hESC line). Data normalized to engulfment in the mock control. Student’s *t* test. ** represents ***P < 0.001. Data are presented as mean ± s.e.m.

Abnormalities in synaptic pruning have been reported in many neurological disorders, brain injury, and viral infection (Garber et al., 2019; Gunner et al., 2019; Hong et al., 2016; Sellgren et al., 2019). To examine whether ZIKV exposure alters human microglial synaptic pruning function, we infected day 75 organoids, in which synapses have formed. As shown in Figures 6I and 6J, compared with mock-infected control, ZIKV exposure significantly increased PSD95^+^ engulfment in CD68^+^ microglial phagolysosome compartments, indicating increased microglial synaptic elimination following ZIKV infection. We further examined complement C3, complement receptor 3 (CR3) and the fractalkine receptor (CX3CR1), which mediate major molecular pathways modulating microglia synaptic pruning in development and disease (Bialas and Stevens, 2013; Gunner et al., 2019; Hong et al., 2016). The qRT-PCR results showed that the expression of C3, CR3, but not CX3CR1, increased after ZIKV exposure, suggesting complement cascade, particularly C3 and microglial complement receptor CR3 might be involved in ZIKV-induced synaptic removal (Figure 6I). Taken together, these results show that microglia are able to respond to ZIKV exposure and exhibit increased synaptic elimination in organoids. Thus, this microglia-containing brain organoid model may provide a new platform to study human microglia function in ZIKV infection that was ignored previously due to the lack of this crucial cell type in the organoids.

## Discussion

In this study, we develop a microglia-containing brain organoid model by co-culturing hPSC-derived pNPCs and PMPs at the onset of 3D organoid formation. In this organoid model, PMPs mature into functional microglia, as indicated by their ability to phagocytize NPCs and apoptotic cells as well as prune synapses. Meanwhile, NPCs differentiate into mature neurons that show electrophysiological activities in addition to their complex morphology. Importantly, using this organoid model, we find that human microglia excessively prune synapses under ZIKV infection.

Different from previous microglia-containing brain organoid models (Abud et al., 2017; Ormel et al., 2018), our model has several unique advantages. First, the ratio of microglia can be adjusted by controlling the starting number of pNPCs and PMPs. In the CNS, microglia account for around 5-10% of total CNS cells (Frost and Schafer, 2016). In our study, using a ratio of 7:3 for NPCs to PMPs, we generated organoids with around 8% microglia of the total cell population. Thus, this method can generate organoids with a consistent and physiologically relevant percentage of microglia. Second, since pNPCs are highly responsive to instructive neural patterning cues *in vitro* (Elkabetz et al., 2008; Li et al., 2011), these organoids can be assigned brain region identities at early stages of organoid culture by giving different patterning morphogens. This enables us to study microglia function in a brain region-dependent manner. In this study, we patterned organoid into dorsal or ventral forebrain identity and showed that microglia phagocytized dorsal or ventral forebrain NPCs in the corresponding organoids (Figures 4C, 4D, 4G and 4H). Third, co-culture of fate-committed ectoderm-derived pNPCs and mesoderm-derived PMPs at the onset of organoid formation better mimics early neurodevelopmental processes observed in *in vivo* brain development. This allows NPCs and PMPs to interact and mature reciprocally and significantly reduces possible contamination by other non-neural endoderm- or mesoderm-derived cells. In our day 27 organoids, the expression of *IGF1* and *IL-1β* significantly increased in organoids containing microglia than organoids without microglia (Figure S1D), suggesting that microglia may support neurogenesis by providing these factors in organoids (Casano and Peri, 2015; Shigemoto-Mogami et al., 2014; Ueno et al., 2013). In addition to the protocol we used, several protocols for generating microglia progenitor-like cells from human PSCs have been reported (Douvaras et al., 2017; Guttikonda et al., 2021; Haenseler et al., 2017; McQuade et al., 2018; Muffat et al., 2016). For example, Douvaras et al. demonstrated the generation of monocytes from hPSCs and then differentiation of these monocytes to microglia progenitor-like cells *in vitro*. Human PSCs-derived microglial progenitor cells derived from a different developmental trajectory could be similarly used to make microglia-containing cerebral organoids using the current method.

Our new microglia-containing brain organoid model provides unique opportunities to investigate human microglial function in development and disease, particularly with regard to microglial synaptic pruning function. During CNS development, microglia-mediated synaptic pruning plays a critical role in removing unwanted synapses and shaping neural circuits (Bialas and Stevens, 2013; Hong et al., 2016; Sakai, 2020). A previous study showed that human microglia engulfed post-synaptic materials in organoids, suggesting human microglia might prune synapses in organoids (Ormel et al., 2018). In this study, using super-resolution imaging techniques, we clearly demonstrated that synapses labeled by both pre-synaptic and post-synaptic markers were engulfed in microglia lysosomal compartments in the organoids (Figures 5D, 5E, S3B, and S3C). Different from previous studies which reported either pre- or post-synaptic materials were engulfed by microglia(Ormel et al., 2018; Weinhard et al., 2018), our super-resolution imaging results reveal that human microglia appear to phagocytize the whole synapses. However, the quantification result shows a fewer number of colocalized PSD95^+^/synapsin 1^+^ puncta than single synapsin 1^+^ puncta or PSD95^+^ puncta inside CD68^+^ microglia in day 55 organoids (Figure S3C, S3D). Since neurons were able to form functional synapses and robustly exhibit spontaneous post-synaptic currents in day 90 organoids (Figures 3H, 3J), the lower percentage of the juxtaposition of the puncta complex in day 55 organoids might be due to the immaturity of the neurons at this stage. Moreover, we show that in addition to synaptic pruning, microglia phagocytize NPCs and remove dead cells in the organoid (Figures 4C-4J), recapitulating microglial functions observed in the developing brain(Cunningham et al., 2013; Takahashi et al., 2005). Therefore, our microglia-containing organoid model enables us to study human microglial function in brain development, as well as neurodevelopmental disorders.

Previous studies using brain organoid models have shown that organoids are useful for studying ZIKV mediated pathology in humans. It was previously shown that ZIKV infection causes a reduction in human NPC pools and limits their proliferation in organoids (Dang et al., 2016; Qian et al., 2016). However, these organoids may not be able to fully model the disease mechanisms. For example, some receptors that are crucial for ZIKV infection are expressed by human astrocytes and microglia, but not human NPCs (Di Lullo and Kriegstein, 2017; Meertens et al., 2017; Qian et al., 2016). Thus, it is of particular importance to employ organoids containing diverse types of neural cells for studying ZIKV infection. Using our microglia-containing brain organoid model, we demonstrate that in response to ZIKV infection, human microglia target neurons and excessively remove synaptic materials. In addition to infecting the fetal brain, increasing evidence shows that ZIKV is also associated with a broad range of neurological disorders in adults (da Silva et al., 2017; Figueiredo et al., 2019). Notably, ZIKV can replicate in ex vivo adult human brain tissue (Figueiredo et al., 2019). In our study, after ZIKV infection, we found that microglia adopted a more activated amoeboid-like morphology and expressed elevated levels of inflammatory cytokines. Interestingly, human microglia excessively removed synaptic materials in our model, consistent with previous findings in mice that mouse microglia engulf hippocampal presynaptic terminals following ZIKV acute infection(Figueiredo et al., 2019; Garber et al., 2019). In addition, we also examined C3, a complement protein, and two receptors, CX3CR1 and C3R, which are important for microglia-mediated synaptic pruning during development and disease(Bialas and Stevens, 2013; Gunner et al., 2019; Hong et al., 2016). Consistent with the previous findings that C3 expression was increased in the microglia after ZIKA infection in the mouse brain(Figueiredo et al., 2019), we found increased expression of C3 and C3R after ZIKV infection in the human brain organoid. Furthermore, we also found that expression of INFAR1/2, which play critical roles in antiviral immune responses(Teijaro, 2016), were also upregulated in the brain organoids. Thus, this microglia-containing organoid can be employed to study interactions between human microglia and neurons in response to ZIKV infection and subsequent antiviral immune responses. Moreover, we explored astroglial function, and potential glia-glia and glia-neuron interactions in response to ZIKV infection in our microglia-containing organoids. First, we found the percentage of GFAP^+^ cells in organoids significantly increased after ZIKV exposure compared with mock-infected control (Figures S4A and S4B), consistent with previous studies (Buttner et al., 2019; Huang et al., 2018). In addition, previous studies demonstrate that TNF-α and C3 mediate the cross-talk between astrocytes and microglia (Guttikonda et al., 2021). For example, in a recent study (Guttikonda et al., 2021), Guttikonda et al. found a neuroinflammatory loop between microglia and astrocytes in response to LPS stimuli. In this loop, LPS initially activated microglia to increase the production of C3. Next, stimulated microglia signal to astrocytes by releasing TNF-α, and astrocytes in turn signal back to microglia by C3 secretion. In our study, increased expression of *TNF-α* and *C3* was observed in organoids after ZIKA infection, which might implicate potential interaction between astrocyte and microglia. It would be interesting in future studies to further explore whether a similar loop exists in ZIKA infection and whether TNF-α and C3 derived from astrocytes or microglia could further impact neuronal functions.

Increasing evidence indicates that adaptive immune cells play essential roles in brain development and disease (Pasciuto et al., 2020; Prinz and Priller, 2017). Future studies incorporating adaptive immune cells in brain organoid cultures will further present new opportunities for studying interactions between human CNS and the adaptive immune system in development and neurological disorders, including virus infection-associated brain dysfunctions such as ZIKV and SARS-CoV2 infections (Ellul et al., 2020; Iadecola et al., 2020). Overall, our new microglia-containing organoid model will provide unique opportunities to investigate human microglial function in development and disease, as well as provide a platform for drug screening.

## Methods

### Generation, culture, and quality control of hPSC lines

Two healthy control hiPSC lines (female and male) and the female H9 ESC line were used in this study. The hiPSC lines were generated from healthy adult-derived fibroblasts using the “Yamanaka” reprogramming factors in our previous study (Chen et al., 2014). The hPSC line has been fully characterized, including karyotyping, teratoma assay, Pluritest (http://www.PluriTest.org), DNA fingerprinting STR (short tandem repeat) analysis and gene expression profiling(Xu et al., 2019; Xu et al., 2020). The hPSCs, from passage number 15–30, were cultured on dishes coated with hESC-qualified Matrigel (Corning) in mTeSR1 medium (STEM CELL Technologies) under a feeder-free condition and were passaged once per week with ReLeSR medium (STEM CELL Technologies). All hPSC studies were approved by the Embryonic Stem Cell Research Oversight committee at Rutgers University.

### pNPC generation and culture

Human pNPCs were generated from hPSCs as previously reported(Chen et al., 2016; Xu et al., 2019). Briefly, to induce neural differentiation, embryoid bodies (EB) were treated with noggin (50 ng/ml, Peprotech) and SB431542 (5 mM, Stemgent) for dual inhibition of SMAD signaling, in neural induction medium consisting of DMEM/F12 (HyClone) and 1x N2 (Thermo Fisher Scientific) for 1 week. EBs were then plated on dishes coated with growth factor-reduced Matrigel (BD Biosciences) in medium composed of DMEM/F12, 1x N2, and laminin (1 mg/ml; Corning). pNPCs emerged in the form of neural rosettes for another 7 days, after which, rosettes were manually isolated from surrounding cells and expanded for another 7 days in pNPC medium, which is composed of a 1:1 mixture of Neurobasal (Thermo Fisher Scientific) and DMEM/F12, supplemented with 1x N2, 1x B27-RA (Thermo Fisher Scientific), FGF2 (20 ng/ml, Peprotech), CHIR99021 (3 mM, Biogems), human leukemia inhibitory factor (hLIF, 10 ng/ml, Millipore), SB431542 (2 mM), and ROCK inhibitor Y-27632 (10 mM, Tocris). Next, the expanded pNPCs were dissociated into single cells using TrypLE Express (Thermo Fisher Scientific) for organoid generation or long-term storage in liquid nitrogen.

### PMP generation and culture

The PMPs were derived from hPSCs, as previously reported(Haenseler et al., 2017; Xu et al., 2020), and produced in a Myb-independent manner to recapitulate primitive hematopoiesis(Buchrieser et al., 2017; Haenseler et al., 2017). Briefly, yolk sac embryoid bodies (YS-EBs) were generated by treating EBs with bone morphogenetic protein 4 (BMP4, 50 ng/ml, Peprotech; to induce mesoderm), stem cell factor (SCF, 20 ng/ml, Peprotech), and vascular endothelial growth factor (VEGF, 50 ng/ml, Peprotech) in mTeSR1 medium (STEM CELL Technologies). Then, the YS-EBs were plated into dishes and cultured in PMP medium, which is composed of X-VIVO 15 medium (Lonza), supplemented with macrophage colony-stimulating factor (M-CSF,100 ng/ml, Peprotech) and interleukin-3 (IL-3, 25 ng/ml, Peprotech) to promote myeloid differentiation. After 2–3 weeks of plating, human PMPs begin to emerge into the supernatant and are continuously produced for around 3 months. The cumulative yield of PMPs can reach around 40-fold of input hPSCs.

### Brain organoid generation and culture

To generate brain organoids, 10,000 cells were used per organoid similar to our previous studies (Kim et al., 2019; Xu et al., 2019). In this study, 7,000 pNPCs plus 3,000 PMPs were co-cultured, supplemented with ROCK inhibitor Y-27632 (10 mM), in ultra-low-attachment 96-well plates to form uniform organoids in medium composed of a 1:1 mixture of PMP medium and NPC medium. NPC medium is composed of a 1:1 mixture of Neurobasal (Thermo Fisher Scientific) and DMEM/F12, supplemented with 1x N2, 1x B27-RA (Thermo Fisher Scientific), FGF2 (20 ng/ml, Peprotech) for three days (day 24). Then, organoids were transferred to ultra-low-attachment 6-well plates and cultured with 1:1 mixture of PMP medium and NPC medium for another 11 days (day 35). To pattern these organoids to the fate of dorsal or ventral forebrain, we respectively culture the organoids without morphogen or treated organoids with sonic hedgehog (SHH) (50 ng/mL, Peprotech) and purmorphamine (1 mM, Cayman Chem), an agonist of sonic hedgehog signal pathway from day 21 to day 35(Figure 1A). The medium was replenished every day, and the cell culture plates were kept on the orbital shaker with a speed of 80 rpm/min starting from day 27. Maximum 8 organoids were cultured in each well of the ultra-low-attachment 6-well plate. To promote neural and microglial differentiation, organoids were cultured in differentiation medium, which contained a 1:1 mixture of Neurobasal and DMEM/F12, supplemented with 1x N2 (Thermo Fisher Scientific), BDNF (20 ng/ml, Peprotech), GDNF (20 ng/ml, Peprotech), dibutyryl-cyclic AMP (1mM, Sigma), ascorbic acid (200 nM, Sigma), IL-34 (100 ng/ml, Peprotech), Granulocyte-macrophage colony-stimulating factor (GM-CSF, 10 ng/ml, Peprotech) from day 35 onwards. Medium was replenished every other day. To further promote neural maturation, organoids were cultured in media comprised of 1:1 mixture of the aforementioned differentiation media and BrainPhys neuronal medium (STEMCELL Technologies). These organoids were used for functional patch-clamp recordings.

### Immunostaining and cell counting

Immunostaining and cell counting were performed similarly to our previous report(Xu et al., 2019). Briefly, organoids were fixed with 4% paraformaldehyde (PFA) for 1h at room temperature and then immersed in 25% sucrose at 4 °C overnight for cryoprotection. Organoids were then embedded in OCT medium, snap frozen in a dry-ice chilled methanol bath and sectioned in a cryostat (Leica) at 18-μm thickness (unless otherwise specified). For immunofluorescence staining, the sections were washed with PBS before permeabilization/blocking with 5% goat or donkey serum in PBS with 0.2% Triton X-100 at room temperature (RT) for 1 h. The sections were incubated with primary antibodies diluted in the same blocking solution at 4 °C overnight. All antibodies used are listed in Supplementary Table 1. Then, the sections were washed with PBS and incubated with secondary antibodies for 1 h at RT. Finally, the slides were washed with PBS and mounted with anti-fade Fluoromount-G medium containing 1, 40,6-diamidino-2-phenylindole dihydrochloride (DAPI) (Southern Biotechnology). A Zeiss 710 confocal microscope was used for capturing images. Zeiss Airyscan super-resolution microscope at 63X with 0.2 mm z-steps was used to acquire super-resolution images for visualizing synaptic puncta pruning. Imaris software (Bitplane 9.5) was used to process super-resolution images and generate 3D-surface rendered images as previously reported(Xu et al., 2020). To quantify the endpoint number and process length of microglia in organoids, the Filament function of Imaris software was employed. To quantify the density and percentage of synapsin 1^+^, PSD95^+^, and colocalized PSD95^+^ puncta (PSD95^+^ puncta in the juxtaposition of puncta complexes) along dendrites, we calculate the number of puncta per 10 μm^2^ MAP2^+^ dendrites using ImageJ as previously reported (Catarino et al., 2013; Patzke et al., 2019) (Ciani et al., 2011). To visualize the engulfed puncta within microglia, the mask function of Imaris was used to subtract any fluorescence that was outside of the microglia, as previously reported(Schafer et al., 2014). The cells were counted with ImageJ software, and at least three fields of each organoid were chosen randomly for cell counting. To quantify synapsin 1, PSD95, and juxtaposition of puncta within the phagosome, the number and volume of synaptic puncta and CD68^+^ phagosome (Figures 5F, S3D and 6J), Imaris was utilized as previously reported (Gunner et al., 2019; Schafer et al., 2014). In ZIKA infection (Figure 6J), all data were then normalized to the control as previously reported (Schafer et al., 2014). For the visualization of neurons filled with Neurobiotin (Biotin Ethylenediamine HBR) during patch-clamp recording, slices were immediately fixed for 30 min at RT in 4% PFA. Slices were washed with PBS, permeabilized for 45 min in the same blocking solution above, and incubated with Alexa Fluor 546 streptavidin conjugate in blocking solution for 1 hr at RT. Finally, slices were washed with PBS and mounted in Fluoromount-G medium for subsequent confocal imaging.

### RNA isolation and qPCR

Total RNA was extracted from organoids using Trizol (Invitrogen), as previously reported(Xu et al., 2014). Complementary DNA was prepared using a Superscript III First-Strand kit (Invitrogen). qPCR was performed with Taqman primers (Life Technologies) on an ABI 7500 Real-Time PCR System. All primers used are listed in Supplementary Table 2. All experimental samples were analyzed and normalized with the expression level of housekeeping gene glyceraldehyde-3-phosphate dehydrogenase (GAPDH). Relative quantification was performed by applying the 2-DDCt method(Xu et al., 2019).

### Whole-cell patch-clamp recording in organoid slices

Organoids were embedded in low melting temperature agarose (SeaPlaqueTM Low Melt Agarose) and sectioned into 200-300 μm thick sections in ice-cold oxygenated (95% O_2_/5% CO_2_) ACSF containing the following (in mM): 125 NaCl, 2.5 KCl, 1.25 NaH_2_PO_4_, 25 NaHCO_3_, 2.5 glucose, 50 sucrose, 0.625 CaCl_2_, and 1.2 MgCl_2_. The osmolarity of the cutting solution was 320-330 mOsm. The slices were then transferred to a recovery bath for 45 min at 32°C in oxygenated (95% O_2_/5% CO_2_) ACSF containing the following (in mM): 125 NaCl, 2.5 KCl, 1.25 NaH_2_PO_4_, 25 NaHCO_3_, 2.5 glucose, 22.5 sucrose, 2.5 CaCl_2_, and 1.2 MgCl_2_. The osmolarity of the external solution was 305-315 mOsm. Electrophysiological recordings were performed in the same solution with a perfusion rate of 2 ml/min at room temperature. Both cutting and patching external ACSF solutions had a pH 7.2-7.4 when saturated with 95% O_2_/5% CO_2_. Neurons were identified by differential interference contrast (DIC) optics under an upright microscope (bx51, Olympus) with a 40x water-immersion objective. Patch pipettes were pulled from borosilicate glass with a resistance of 5-8 mΩ. For current-clamp and voltage-clamp recordings, a K-gluconate internal solution was used consisting of (in mM): 126 K-gluconate, 4 KCl, 10 HEPES, 0.05 EGTA, 4 ATP-magnesium, 0.3 GTP-sodium, and 10 phosphocreatine. The pH was adjusted to 7.2 with KOH and the osmolarity adjusted to 280-290 mOsm. Neurons were held at −70mV for voltage-clamp recordings. In combination with the electrophysiological recordings, we added 0.5% Biotin Ethylenediamine HBR (Biotum) to the internal solution in order to label patched neurons. Whole-cell patch-clamp recordings were performed using an Axon 700B amplifier. Data were filtered at 2 kHz, digitized at 10 kHz, and collected using Clampex 10.6 (Molecular Devices). Electrophysiological data were analyzed offline using Clampfit 10.6 software (Molecular Devices).

### Zika virus infection

Day 75 old organoids were infected with ZIKV. Two different strains were tested initially, including MR766 (Uganda) and Fortaleza (Brazil), and MR766 was used for most experiments. Briefly, organoids were infected with ZIKA or mock, with a concentration of 10^4^ PFU/ml in 6 well plates on the orbital shaker. After three days of infection, organoids were replenished with fresh medium without ZIKV and cultured for another four days. At day three or day seven post-infection, organoids were collected for immunohistochemistry or RNA extraction, and the supernatant was collected for plaque assay. Briefly, Vero CCL-81 cells (ATCC) were plated at 2E5 cells/well in 12-well plates. Supernatant from infected organoids was serially diluted 10-fold, and 100μL transferred to each well. Plates were rocked every 15 minutes for 1 hour before a semi-solid overly containing 0.5% Methylcellulose in viral growth media (VWR) was applied. 4 days post infection, the overlay was removed, and cells fixed in a solution of 10% neutral buffered formalin with 0.25% crystal violet for plaque counting.

### Statistics and reproducibility

All data represent mean ± s.e.m. When only two independent groups were compared, significance was determined by two-tailed unpaired t test with Welch’s correction. A p-value <0.05 was considered significant. The analyses were done in GraphPad Prism v.5. All experiments were independently performed at least three times with similar results.

## Acknowledgements

This work was in part supported by grants from the NIH (R21HD091512, R01NS102382, and R01NS122108 to P.J.). A.J.B is supported by the National Institute of General Medicine Sciences (NIGMS) NIH T32 GM008339.

## Author Contributions

P.J. and R.X. designed experiments and interpreted data; R.X. carried out most of experiments with technical assistance from X.L., C.E., and M.J.; A.J.B. and Z.P. performed patch-clamp recordings. C.A, and B.P.D. performed ZIKV infection experiments. Z. P. and B.P.D. provided critical suggestions. P.J. and R.X wrote the manuscript with input from all co-authors.

---

**Figure S1.**
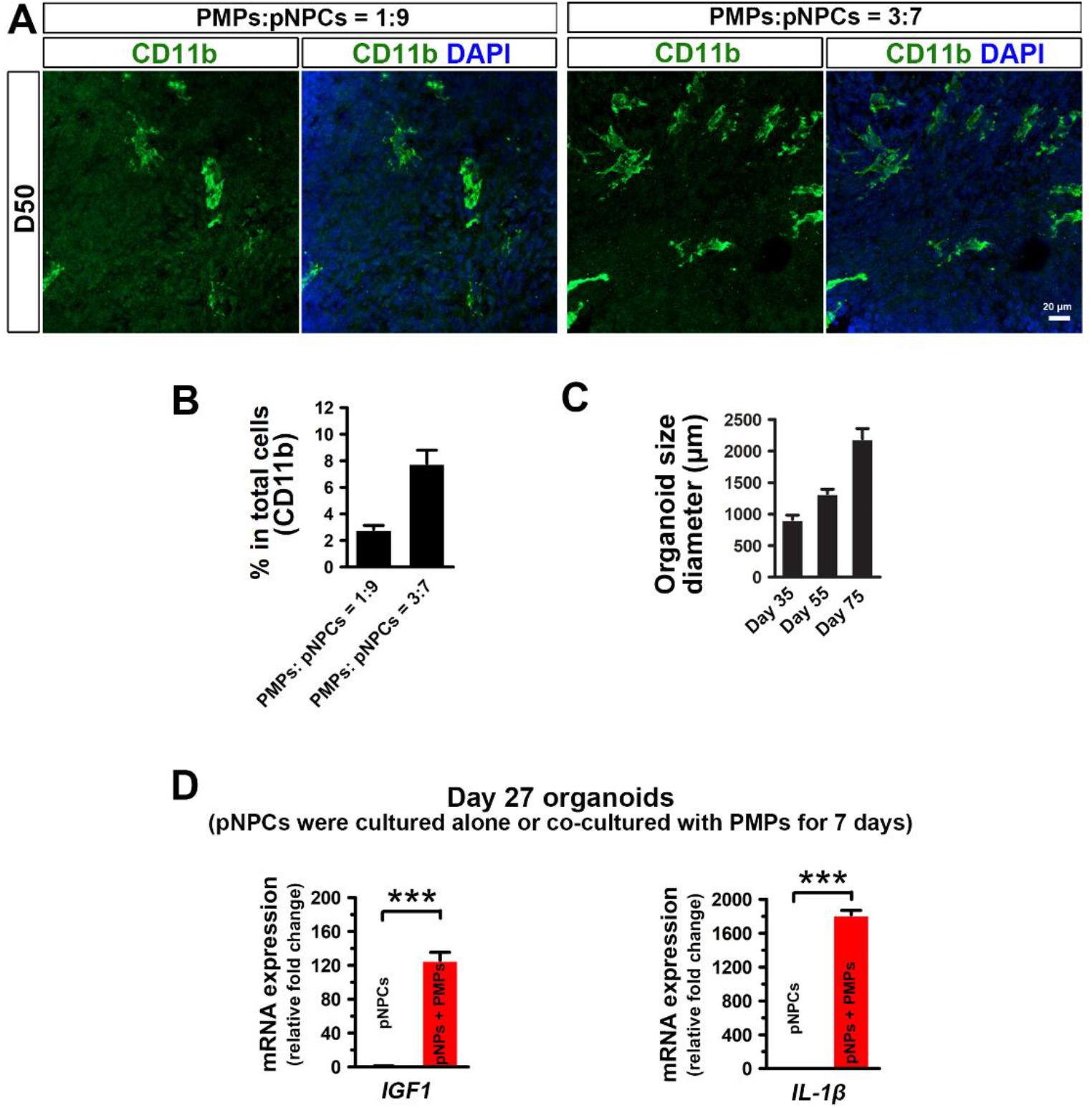
Characterization of microglia-containing organoids. (A) Representatives of CD11b^+^ cells in day 50 brain organoids with different starting numbers of PMPs and pNPCs. Scale bars: 20 μm. (B) Quantification of pooled data from an hiPSC and an hESC line showing the percentage of CD11b^+^ cells in day 50 organoids with different starting numbers of PMPs and pNPCs (n = 3). Three independent experiments were performed, and for each experiment, 4-6 organoids from each hPSC line were used. Data are presented as mean ± s.e.m. (C) Quantification of organoid size at different stages. Scale bars: 200 μm. Data are presented as mean ± s.e.m. (D) qPCR analysis of *IGF1* and *IL-1β* mRNA expression in day 27 organoids (co-culture for 7 days) with or without microglia (n = 3, each time, 30-40 organoids pooled from two hiPSC lines). Student’s *t* test. *** represents ***P < 0.001. Data are presented as mean ± s.e.m.

**Figure S2.**
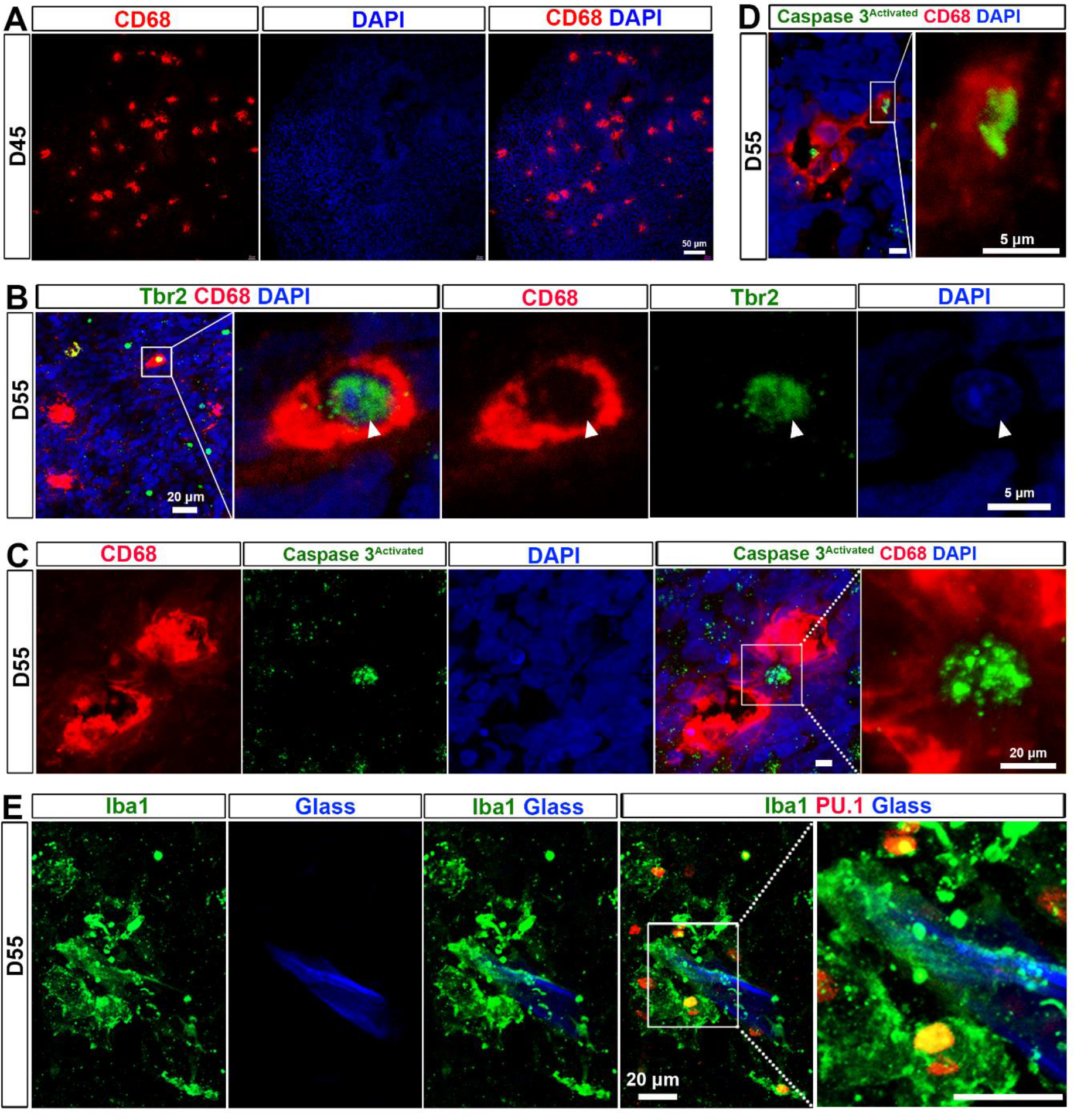
Phagocytic function of microglia in brain organoids. (A) Representative of CD68^+^ cells in day 45 organoids. Scale bars: 50 μm. (B) Representative images showing CD68^+^ microglia phagocytosing Tbr2^+^ intermediate progenitor cells in day 55 organoids. The enlarged images. Scale bars: 20 μm or 5 μm in the original or enlarged images, respectively. (C and D) Representative images showing CD68^+^ microglia phagocytosing activated-Caspase-3^+^ cell in day 55 organoids. Scale bars: 20 μm. (E) Representative images showing Iba1^+^ and PU.1^+^ microglia accumulate around foreign body glass fragments from pipette tip in day 55 organoids. Scale bars: 20 μm.

**Figure S3.**
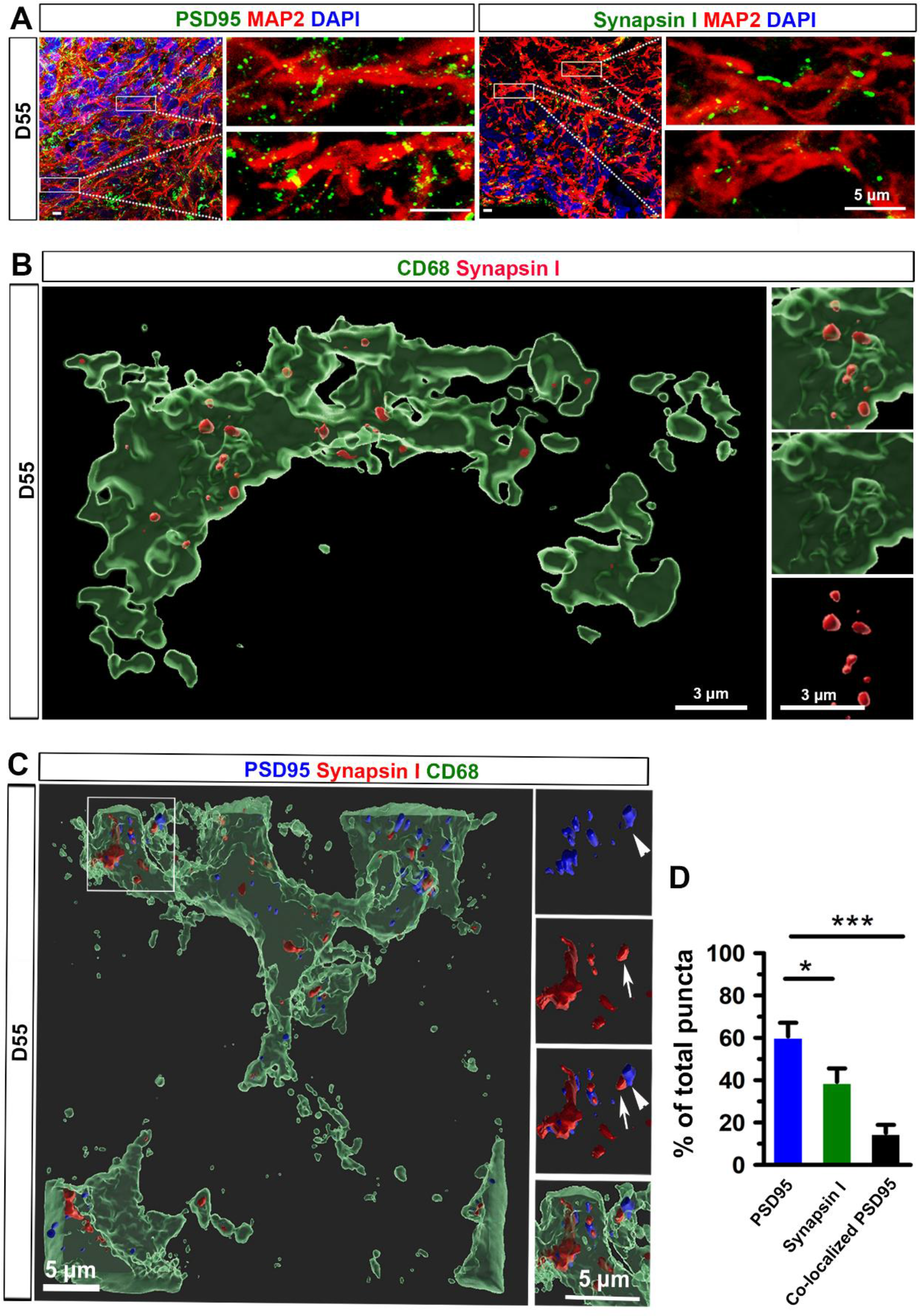
Synaptic pruning function of microglia in brain organoids. (A) Representative images showing PSD95^+^ and synapsin 1^+^ synaptic puncta along MAP2^+^ processes in day 55 brain organoids. Scale bars: 20 μm or 5 μm in the original or enlarged images, respectively. (B) Representative 3D reconstruction of super-resolution images showing synapsin 1^+^ puncta within CD68^+^ microglial phagolysosomes in day 55 organoids. Scale bars, 3 μm. (C) Representative 3D-surface rendered images showing CD68^+^ microglial phagolysosomes engulfment of synapsin 1^+^ and PSD95^+^ puncta in day 55 organoids. Arrows and arrowheads indicate the juxtaposition of pre- and postsynaptic puncta, respectively. Scale bars, 5 μm. (D) Quantification of pooled data from two hiPSC lines and one hESC line showing the percentage of synapsin 1^+^, PSD95^+^, and colocalized PSD95^+^ per 100 μm^3^ CD68^+^ microglial phagolysosomes in day 55 organoids (n = 3, three lines each time, 4-6 organoids each line each time). One-way ANOVA test. *p < 0.05 and ***p < 0.001. Data are presented as mean ± s.e.m.

**Figure S4.**
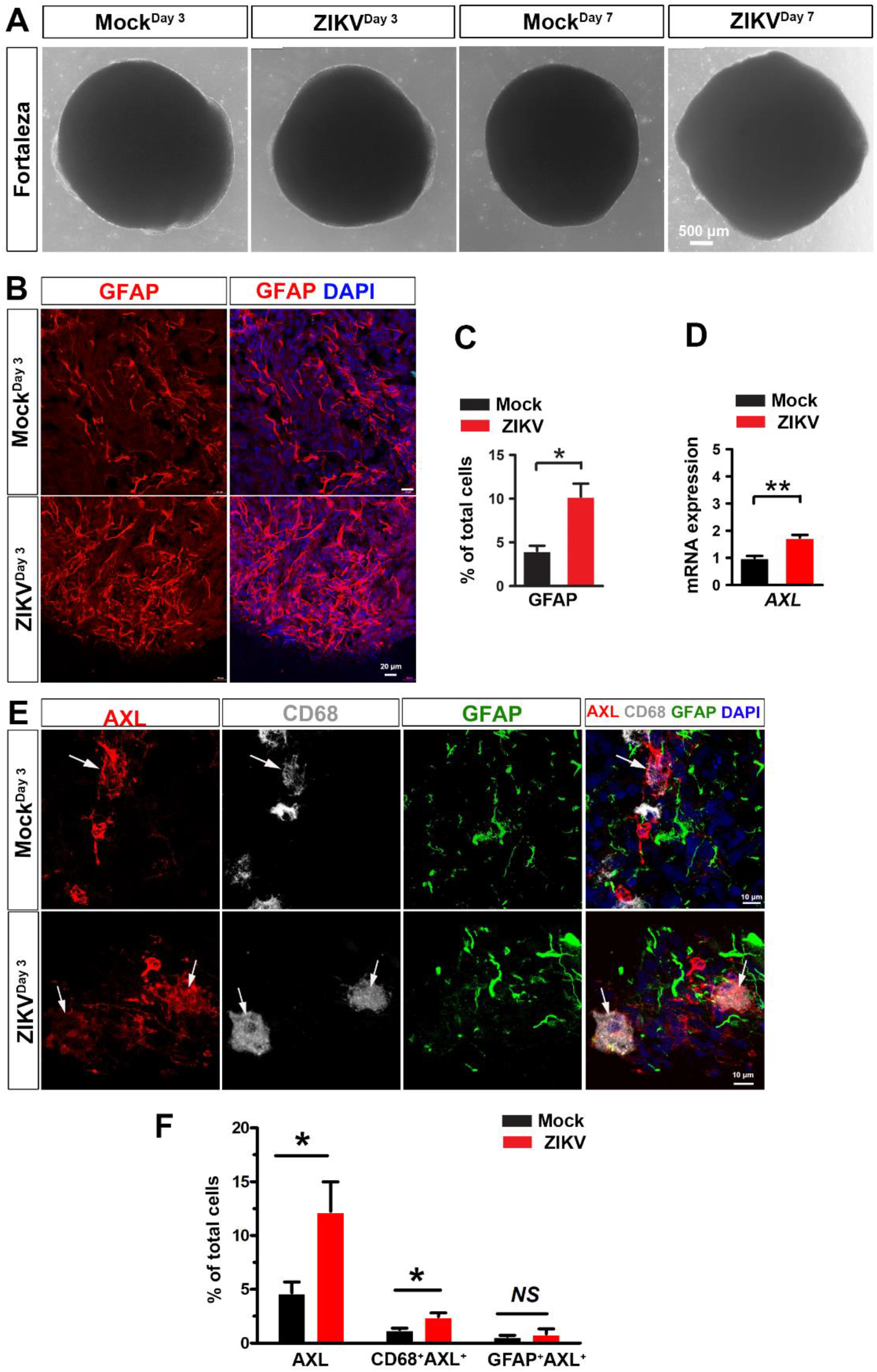
Microglia-containing organoids respond to ZIKV infection. (A) Representative phase-contrast images showing the morphology of day 75 organoid at 3- and 7-day day post Fortaleza ZIKV-Fortaleza infection. Scale bars: 500 μm. (B) Representative images showing GFAP^+^ astrocytes in day 75 organoids after three days of ZIKV (MR766) or mock infection. Scale bars: 20 μm. (C) Quantification of the percentage of GFAP^+^ astrocytes in day 75 organoids after three days of ZIKV or mock infection. (n = 3, three times of independent infections; each time, 9-12 organoids from two hiPSC lines and one hESC line). Student’s t test. *P < 0.05. Data are presented as mean ± s.e.m. (D) qPCR analysis of *AXL* mRNA expression in day 75 organoid after three days of ZIKV or mock infection. (n = 3, three times of independent infections; each time, 8-12 organoids from two hiPSC lines). Student’s *t* test. **P < 0.01. Data are presented as mean ± s.e.m. (E) Representative images showing AXL^+^, CD68^+^, and GFAP^+^ cells in day 75 organoids (three days after MR766 ZIKV or mock infection). Arrows indicate CD68^+^/AXL^+^ microglia. Scale bars: 10 μm. (F) Quantification of the percentage of AXL^+^ cells, CD68^+^/AXL^+^ microglia, and GFAP^+^/AXL^+^ astrocytes in day 75 organoids. (n = 4-6 organoids from one hiPSC line). Student’s t test. *P < 0.05. Data are presented as mean ± s.e.m.

